# Role of long-range aromatic clique and community in protein stability

**DOI:** 10.1101/319160

**Authors:** P Mahanta, A Bhardwaj, VS Reddy, S Ramakumar

## Abstract

Aromatic interactions make an important contribution to protein structure, function, folding and have attracted intense study. Earlier studies on a recombinant xylanase from Bacillus sp. NG-27 (RBSX), which has the ubiquitous (beta/alpha)_8_-triosephosphate isomerase barrel fold showed that three aromatic residues to alanine substitutions, in the N-terminal and C-terminal regions, significantly decreased the stability of the enzyme. Of these mutations, F4A mutation decreased the stability of the enzyme by ∼4 degree C, whereas W6A mutation and Y343A mutation remarkably decreased the stability of the enzyme by ∼10 degree C. On the other hand, the F4W mutation did not affect the thermal stability of RBSX. We provide here a network perspective of aromatic-aromatic interactions in terms of aromatic clique community and long-range association. Our study reveals that disruption of long-range k-clique aromatic interaction cluster holding the N- and C-terminal regions are associated with the decreased stability of the enzyme. The present work reiterates as well as expands on those findings concerning the role of interactions between the N- and C-terminus in protein stability. Furthermore, comparative analyses of crystal structures of homologous pairs of proteins from thermophilic and mesophilic organisms emphasize the prevalence of long-range k-clique communities of aromatic interaction that may be playing an important role and highlights an additional source of stability in thermophilic proteins. The design principle based on clustering of long-range aromatic residues in the form of aromatic-clique and clique community may be effectively applied to enhance the stability of enzymes for biotechnological applications.

**Database:** The coordinates o fF4A, F4W, W6A, and Y343A are deposited in the PDB database under the accession numbers **5EFF**, **5E58**, **5EFD**, and **5EBA** respectively.

**Abbreviations:** BSX, xylanase from *Bacilllus sp.* NG-27; RBSX, recombinant BSX xylanase; TIM, Triosephosphate isomerase; GH10, Glycosyl hydrolase family 10; 3D, three-dimensional; r.m.s.d, root mean square deviation; RSA, relative solvent accessible surface area; T_m_, melting temperature; CD, Circular Dichroism; BHX, GH10 xylanase from *Bacillus halodurans*; BFX, GH10 xylanase from *Bacillus firmus*; TmxB, GH10 xylanase from *Thermotoga maritima*

## Introduction

Proteins play vital role in cellular processes of organisms. Although proteins are composed of twenty amino acids; irrespective of their source organisms, they adapt to different physico-chemical environments such as low or high temperature, high pressure, high salinity, a broad pH range, or resistance to protease degradation and combinations, leading to poly-extremophilic stability [1]. There are multiple factors governing the stability of a protein; however, each protein appears to have evolved its own mechanism to achieve a high thermal stability rather than converging on a single universal feature [2, 3].

The relevance of aromatic-aromatic interactions has attracted considerable interest in recent times for their key roles in protein structure stability, function, folding, and ligand recognition [4-6]. Both small molecule and macro-molecular crystallographic studies have provided key experimental and theoretical information on aromatic interactions and other non-covalent interactions. Georis et al. (2000) demonstrated that introduction of an additional aromatic interaction (Tyr11-Tyr16) by a single amino acid substitution of T11Y improves the thermostability and thermophilicity of a family 11 xylanase [7]. In an *in silico* study of the homologous pair of thermophilic and mesophilic proteins, Kannan and Vishveshwara (2000) identified that thermophilic proteins have additional or enlarged aromatic clusters than their mesophilic counterparts [6]. In a recent analysis based on the structural information available in the Protein Data Bank (PDB) Lanzarotti et al. (2011) shows the prevalence of aromatic trimers, tetramers and even large clusters and pointed out the relevance of aromatic clusters beyond dimer in protein function, stability, and ligand recognition [8]. However, there is hardly any discussion so far on the role of long-range aromatic interactions and aromatic interactions between protein termini in protein stabilization/destabilization. The present study envisages filling the lacunae and provides a network perspective of aromatic-aromatic interactions in terms of cliques, clique communities and long-range association.

BSX is an extra-cellular endo-xylanase from *Bacillus sp. NG-27*, belongs to glycosyl hydrolases family 10 (GH10) and catalyzes the hydrolysis of internal β-1,4 bonds of xylan backbones [1]. BSX is optimally active at a temperature of ∼70 °C and at a pH 8.5. It has a (β/α)_8_-triosephosphate isomerase (TIM) barrel fold, which is a common tertiary fold, occurring in many glycosyl hydrolases and present in approximately 10% of all enzymes [9]. Earlier studies of recombinant BSX (RBSX) showed that three aromatic residues to alanine substitutions in the N-terminal and C-terminal regions significantly decrease the stability of the enzyme [10]. Of these mutations, F4A mutation decreased the stability of the enzyme ∼by 4 °C, whereas W6A mutation and Y343A mutation remarkably decreased the stability of the enzyme by ∼10 °C. On the other hand, F4W mutation did not affect the thermal stability of RBSX. To provide a structural basis for how aromatic mutations influence the thermal stability of RBSX, we determined the crystal structure of these four mutants. The availability of various experimentally determined aromatic mutant structures has enabled a critical examination of factors influencing thermal stability of the enzyme. Because aromatic residues can form aromatic cluster/s and can be viewed as a network of aromatic residues, network analysis was adopted as a tool to study these interaction networks. We identify an aromatic cluster involving residues Phe4, Trp6 and Tyr343, forming a long-range k=3 k-clique of aromatic interaction network, holding together the N-terminal and C-terminal of RBSX (Fig. 1). Here, a k-clique’ is a fully connected subgraphs of k-nodes in which each node connects with all other nodes in the network. Based on network analysis of aromatic interactions, our study reveals that disruption of long-range aromatic interaction community of k=3 k-clique at the termini is associated with the decreased stability of the enzyme. Furthermore, the present results reiterate and exemplify concerning the role of N-terminal and C-terminal contacts specifically through aromatic interactions or packing in protein stability. Lastly, we carried out a comparative analysis of homologous pairs of proteins from thermophilic and mesophilic organisms and observed in general, higher prevalence of long-range k-clique communities of aromatic interaction in the thermophilic proteins in comparison to their mesophilic homologs, and highlights an additional source of stability in thermophilic proteins. The present study points out that the strategy of mutations based on clustering of aromatic residues in terms of ‘aromatic-clique’ and ‘clique-community’ may be effectively applied to enhance the stability of enzymes and provides useful strategies for the design of proteins for biotechnological applications.

**Fig. 1.**
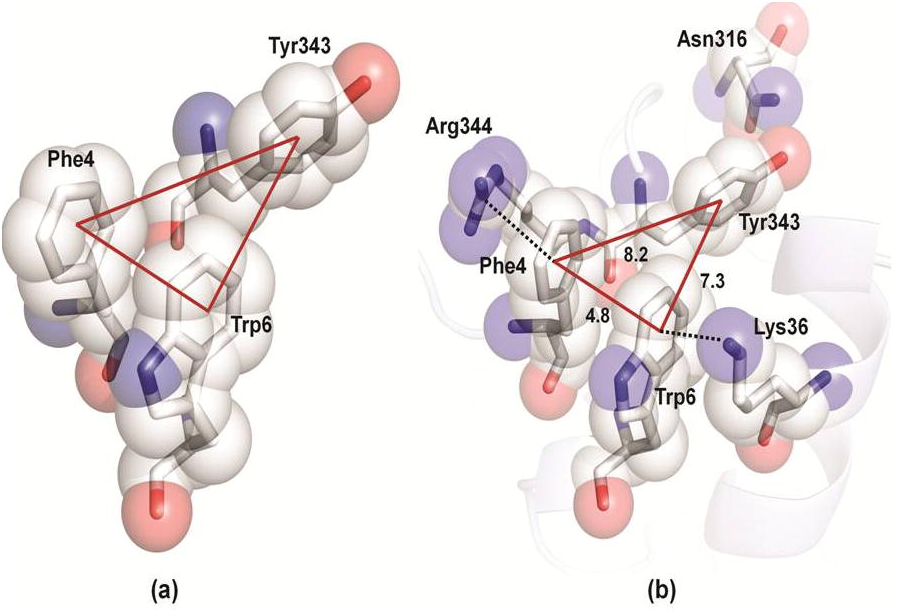
k=3 k-clique long-range aromatic interaction cluster (Phe4-Trp6-Tyr343) of RBSX structure. (a) Example of a k-clique long-range interaction cluster (k = 3) of aromatic residues. (b) Structural feature of k-clique aromatic interaction cluster and their interactions in RBSX structure (PDB ID: 4QCE). The side chains of Phe4, Trp6, Lys36, Asn316, Tyr343, and Arg344 are shown in space filling and stick representation and the backbone is shown as a cartoon. The solid red line represents aromatic interaction whereas dashed blue line represents cation-π interaction.

## Results

### Loss of aromatic interaction destabilizes the native state of RBSX

In an earlier study from our group, we reported a systematic investigation of a set of aromatic mutants of RBSX and showed that substitution with alanine in all these aromatic residues (F4A, F4W, W6A and Y343A) significantly decreases the stability of the enzyme [10]. The thermal stability of RBSX and the mutant proteins were determined by CD measurements at a range of temperatures [10]. RBSX and its mutants were found to unfold in an irreversible manner and the apparent melting temperature (T_m_) was calculated for all protein samples with a constant temperature slope of 60 °C h 1. It was observed that substitution of Phe4 by alanine (F4A) decreased the stability of the enzyme by ∼4 °C, whereas W6A and Y343A mutations significantly decreased the stability by ∼10 °C relative to RBSX. On the other hand, F4W mutation did not alter the stability of the enzyme (Table S1). To gain structural insights into how the mutation influences thermostability of the enzyme, we determined the crystal structure of all four mutants. A brief summary of crystallization, data collection, structure solutions and refinement statistics of different mutants is given in Table 1 and Doc. S1.

**Table 1.**
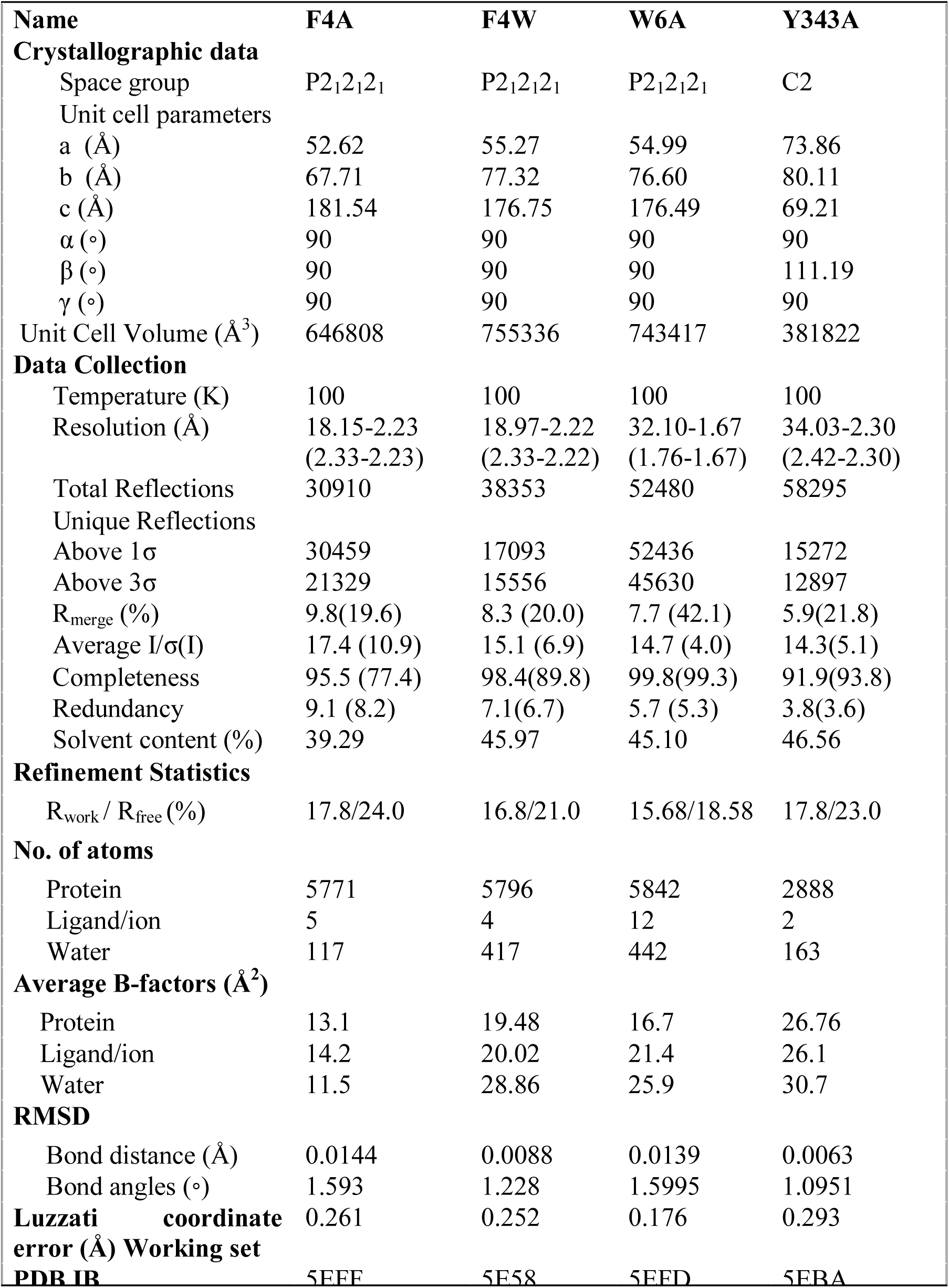
X-ray collection and refinement statistics.

### Structural features of Phe4-Trp6-Tyr343 aromatic cluster in RBSX

RBSX is a monomeric GH10 xylanase having (β/α)_8_-TIM barrel fold [11]. The native structure of RBSX is composed of twelve α-helices, nine β-strands and five 3_10_-helices, as assigned by DSSP [12]. The barrel forming secondary structures consisting of eight major parallel β-strands is lying in the middle, surrounded by eight α-helices. Crystal structure analysis of RBSX reveals an aromatic cluster viz. ‘aromatic-clique’ involving residues Phe4, Trp6, and Tyr343 in which each residue is connected through aromatic interactions to all other residues within the cluster (**Fig. 1b**). The three distances between the center of mass (COM) of the aromatic rings and the three minimum distances between any two atoms belonging to the aromatic ring characterize the existence of an aromatic-clique (Materials and Methods). Here, Phe4 belongs to a loop and Trp6 belongs to part of 3_10_-helix in the N-terminus region whereas Tyr343 belongs to a loop in the C-terminus region of the protein. The COM-to-COM distances are 4.8 Å for Phe4-Trp6 pair, 7.3 Å for Trp6-Tyr343 pair and 8.2 Å for Phe4-Tyr343 pair. On the other hand, the closest distances between any two atoms belonging to the aromatic rings are 3.5 Å (Phe4-Trp6), 4.6 Å (Trp6-Tyr343) and 5.9 Å (Phe4-Tyr343) and the dihedral angles are 10.2° (Phe4-Trp6), 38.9° (Trp6-Tyr343) and 39.1° (Phe4-Tyr343).

### Comparative analysis of different aromatic mutant structures with that of RBSX structure

The crystal structure of F4A, F4W, W6A and Y343A mutants was solved by molecular replacement method at a resolution of 2.23 Å, 2.22 Å, 1.67 Å and 2.30 Å respectively (Table 1,Doc. S1). Comparison of the molecular structures of F4A, F4W, W6A and Y343A mutants with that of RBSX structure showed no significant changes in the overall three-dimensional (3D) structure of proteins. The overall C_α_ r.m.s.d. between RBSX and the F4A mutant, F4W mutant, W6A mutant and Y343A mutant is 0.368Å, 0.189 Å, 0.273 Å and 0.272 Å respectively. The availability of experimentally determined aromatic mutant structures has enabled to examine the local changes at the site of mutation. Evaluation of atomic packing as assessed with the RINERATOR module [13, 14] shows that Phe4, Trp6 and Tyr343 in RBSX structure has a better noncovalent interaction score (*I*_*s*_) [11] (noncovalent *I*_*s*_ is proportional to strength of interaction) than Ala4, Ala6 and Ala343 in the F4A (PDB ID: 5EFF), W6A (PDB ID: 5EFD) and Y343A (PDB ID: 5EBA) mutant structures respectively (detail not shown).

The network analysis of aromatic interactions in RBSX structure reveals that residues Phe4, Trp6 and Tyr343 form a long-range k=3 k-clique aromatic interaction cluster holding the N-terminus and C-terminus together in the 3D space. In the RBSX structure, Phe4 was solvent exposed (relative side-chain solvent accessibility (RSA), 35.9%) and involved in cation-π interactions with Arg344 (Phe4-Arg344) and aromatic interactions with Trp6 (Phe4-Trp6) and Tyr343 (Phe4-Tyr343) (Fig. 1b). The substitution of Phe4 by Ala resulted in the loss of cation-π interaction and aromatic interactions. However, Arg344, which was previously involved in the cation-π interaction with Phe4 in the RBSX structure; now participates in an ionic interaction with Asp340 in F4A mutant structure and seems to be compensating the loss of cation-π interaction. Generally, surface residues of a protein are widely regarded to be tolerant to substitutions. Despite being present at the surface of the protein, the partially exposed Phe4 could modulate the stability of the enzyme. This may be because of the involvement of Phe4 in the long-range N-terminal to C-terminal aromatic cluster network (Phe4-Trp6-Tyr343).

The W6A mutation (located on the N-terminus region) and the Y343A mutation (located on the C-terminus region) results in a substantial decrease in the stability (ΔT_m_ ∼ 10 °C) of the enzyme. In the RBSX structure, the side-chain of Trp6 (RSA= 20.6%) was involved in aromatic interactions with Phe4 (Trp6-Phe4) and Tyr343 (Trp6-Tyr343) and in a cation-π interaction with Lys36 (Trp6-Lys36) (Fig. 1b). The substitution of Trp6 by Ala resulted in the loss of a cation-π interaction and aromatic interactions, thus destabilizing the protein. The hydroxyl group of Tyr343 (RSA = 14.6 %) formed an intra-molecular hydrogen bond to ND2 atom of Asn316 and the side-chain of Tyr343 was involved in aromatic interactions with Phe4 (Phe4-Tyr343) and Trp6 (Trp6-Tyr343) (Fig. 1b). The mutation (Y343A) disrupts the N-terminal to C-terminal aromatic interaction network (Phe4-Trp6-Tyr343) and removes a hydrogen bond between OH of Tyr343 and ND2 of Asn316, thus decreasing the overall stability of the enzyme. Interestingly, substitution with another aromatic residue (F4W) which preserved the N-terminal to C-terminal aromatic interaction cluster (Trp4-Trp6-Tyr343), showed no significant alteration in the stability of the enzyme (Fig. S1). Thus, these observations collectively suggest that the decreased stability shown by the mutants resulted from cumulative effects in the loss of noncovalent interactions (within termini and/or between termini) and primarily by disruption of long-range aromatic cluster between N-terminal to C-terminal of the enzyme. It may be seen that the disruption of long-range aromatic clique community results in destabilization of the enzyme (F4A, W6A, Y343A), whereas no alteration in enzyme stability when the aromatic clique community is maintained (F4W).

### Occurrence of aromatic-clique and community in thermostable GH10 xylanases

It is of interest to investigate whether similar N-terminal to C-terminal aromatic-clique (Phe4-Trp6-Tyr343) is present in other GH10 xylanases. Multiple sequence alignment of RBSX with other GH10 family xylanases from *Bacillus* organisms reveals that aromatic-clique of interest (F-W-Y) is fully conserved in xylanases from *B. halodurans* (BHX) (T_m_ = 70 °C) and *Bacillus firmus* (BFX) (T_m_ = 70 °C), which are thermostable in nature like RBSX (Fig. S2). On the other hand, this particular aromatic-clique of interest is not conserved in GH10 xylanases from *Bacillus N137, Bacillus alcalophilus*; which are reported as thermo-labile in nature. Among the alkaline active (pH 10) thermostable (70 °C) GH10 xylanases produced by an alkalophilic organism, RBSX shows highest sequence similarity score (87.5%) with BHX. Structural superposition of RBSX with BHX (PDB ID: 2UWF) reveals that Phe14, Trp16 and Tyr349 residues form a k=3 k-clique aromatic community in BHX in a similar way to Phe4-Trp6-Tyr343 aromatic-clique community in RBSX and possibly play an important role in their structural stability at elevated temperature (Fig. S3). As the aromatic-clique of interest (Phe4-Trp6-Tyr343) is highly conserved in thermophilic GH10 xylanases from *Bacillus* organisms; to further explore the role of aromatic-clique and community in protein stabilization, we carried out a comprehensive analysis of the thermostable GH10 xylanases with known 3D structures. The analysis shows long-range k=3 k-clique communities of aromatic interaction similar to the one present in RBSX in several members of GH10 thermophilic xylanases (Table S2). For example, two long-range k=3 k-clique aromatic communities have been identified in hyperthermophilic GH10 xylanase TmxB from *Thermotoga maritima* (T_m_ = 90 °C) in which the first aromatic-clique community involves residues Phe532, Phe555, Tyr547, Phe796, Tyr800, Phe816 and Tyr820 whereas the second aromatic-clique community involves residues Tyr727, Tyr766 and Tyr767 respectively. In the first aromatic-clique community of TmxB, Tyr820 (equivalent to Tyr343 in RBSX), belonging to C-terminal region of the protein, participated in aromatic interactions with Tyr547and Phe796 (N-terminal region) and form a long-range k=3 aromatic-clique (Fig. 2). In addition, occurrence of a larger aromatic cluster (Phe532, Phe555, Tyr547, Phe796, Tyr800, Phe816 and Tyr820) involving residues from both N-terminal and C-terminal region in TmxB indicates the association of N-terminal to C-terminal long-range aromatic interaction to protein stability. This feature is also observed in another hyperthermophilic GH10 xylanase from *Thermotoga petrophila RKU-1* (T_m_ = 90 °C) constituting residues from N-terminal and C-terminal regions (Phe35, Tyr50, Phe58, Phe299, Tyr303, Phe319 and Tyr323) (Fig. S4). To surmise, our results show that occurrence of aromatic-clique interactions is a feature common to GH10 thermostable xylanases and might contribute to their stability at higher temperature.

**Fig. 2.**
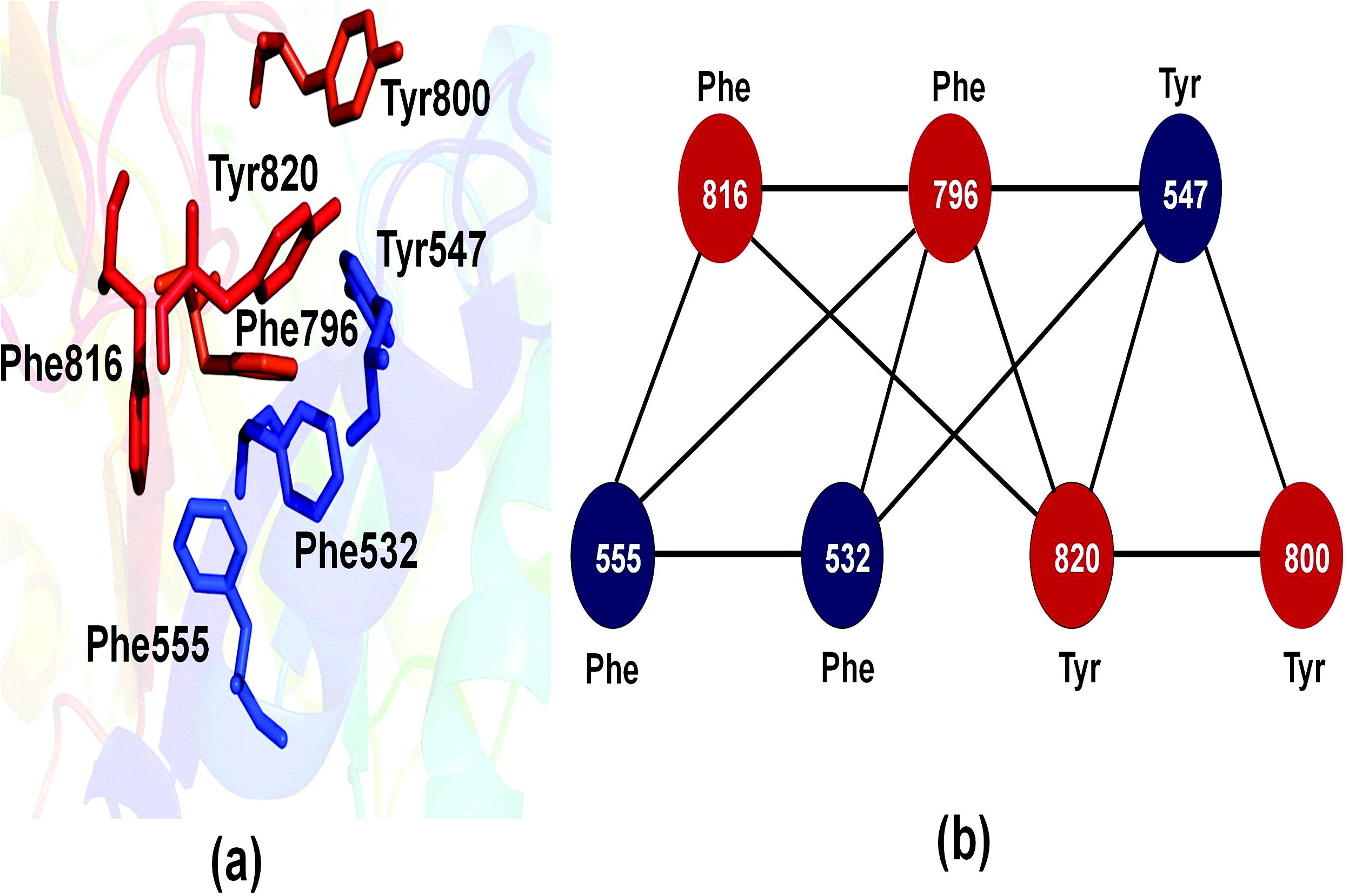
A large cluster of k=3 aromatic-clique community present in hyperthermophilic GH10 xylanases from (a) *Thermotoga maritima* MSB8 (PDB ID: 1VBU, T_m_ = 80 °C). The figure depicts clustering or ‘flocking together’ of aromatic residues in the hyperthermophilic protein. (b) A schematic view of aromatic-clique community present in TmxB GH10 xylanase. It may be seen that three-residue clique is used as a building block to form a large aromatic-cluster.

### Aromatic-clique community: a bioinformatics analysis of homologous pair of proteins from thermophilic and mesophilic organisms

We sought to further generalize our findings by analyzing the occurrence of long-range k=3 k-clique communities of aromatic interaction at structural level in homologous protein pairs from thermophilic and mesophilic organisms. First, we calculated the interacting aromatic pairs and then a k=3 k-clique aromatic community, based on the criteria as described in the materials and methods section. The number of aromatic-clique communities found in the thermophilic proteins and their mesophilic homologs are listed in Table S3. Our analysis reveals preponderance of aromatic-clique communities in thermophilic protein structures, indicating a role for added stability through long-range aromatic interactions that bridge distant regions of the protein. We observed that ∼86% of cases, thermophilic proteins have higher number or equal number of aromatic-clique communities in comparison to their mesophilic homologs. In 37 out of 57 thermophilic-mesophilic pairs, thermophilic proteins have additional long-range aromatic-clique communities which were absent in their mesophilic homologs (Table S3). Here, an additional aromatic-clique community refers to the aromatic-clique community in thermophiles for which topologically equivalent aromatic residues were absent in their mesophilic homologs and vice-versa. We also observed a reverse trend in eight cases (∼14%), in which there is a larger number of additional clique communities in mesophilic proteins than their corresponding thermophilic counterparts. This is presumably because numerous other factors responsible for protein thermostability such as more ion pairs, increase in number of hydrogen bonds, compact packing, and so forth could have been utilized in thermostable proteins. Furthermore, examination of the superimposed structures of the thermophilic and their mesophilic homologs showed that residues forming the additional aromatic-clique communities were mutated to non-aromatic residues in the proteins from mesophiles in which majority of them are hydrophobic residues. For example, an additional long-range aromatic-clique community consisting of residues Tyr77, Tyr78 and Trp138 was identified in the thermophilic xylanase from *Thermomyces lanuginosus* (PDB ID: 1YNA) for which the topologically equivalent residues in the mesophilic protein (PDB ID: 1XYN) are Leu62, Tyr77 and Ile125 (Table S3). Besides, a larger aromatic clique community of five-residues (Tyr14, Tyr77, Trp79, Tyr171 and Tyr172) is observed in the thermophile whereas a smaller aromatic clique community of three residues (Tyr66, Trp68 and Tyr158) is observed in the mesophilic counterpart. Although, Tyr5 is the equivalent aromatic substitution of Tyr14 in the mesophile, it is not part of the aromatic-clique community as Tyr171 which interacts with Tyr14, Trp79 and Tyr172 in the thermophile is mutated to Asn157 in the mesophile. As a result, the interactions are lost and give rise to a smaller cluster (Fig. S5). Another such example comes from the thermophilic RNase H from *thermus thermophiles* (PDB ID: 1JL2), which has an additional long-range aromatic-clique community consisting of residues Phe78, Phe119 and Phe121 for which the topologically equivalent residues in the mesophilic protein are Ile78, Trp118 and Trp120. This feature is observed in ∼65% of the protein families. Thus, the results emerging from the comparative analysis of homologous pairs of protein structures from thermophilic and mesophilic organisms support the idea that aromatic-clique communities are widely utilized by proteins from thermophilic organisms and might be contributing to their thermal stability.

### Three-residue aromatic clique: a building block for larger aromatic cluster

Study of the structural architecture of aromatic clique communities reveals the prevalence of small clusters or small cliques (three residues) which are linked or coalesced to give a large aromatic cluster. For example, a cluster of eleven aromatic residues is observed in Rhodanese from *Thermus Thermophilus HB8* (T_m_ = 70 °C; PDB ID: 1UAR) that comprises of twelve unique three residue cliques (Fig. 3). Another such example comes from Clostridium Thermocellum from *Ruminiclostridium Thermocellum* (PDB ID: 1XYZ) for which a seven residue aromatic cluster is observed that comprises of nine unique three-residue cliques. It emerges that a three residue-clique envisages as the smallest unit of clustering in the protein structure. The three residue clique is of notable importance largely because they could serve as a small building block in the construction of large complex network or could be used to obtain an efficient clustering by growing a seed in the network. The prevalence of higher order aromatic cluster that forms a network of three or more interacting aromatic residues in protein structures was reported by other studies [4, 8]. However, our observation indicates that network of large aromatic clusters could originate from a smaller basis set comprising of three-residue-clique acting as a seed. These clusters have greater link density, highly synergic conformations and reveal its self-associating property to exploit the maximum benefit from clustering in protein stability and function.

**Fig. 3.**
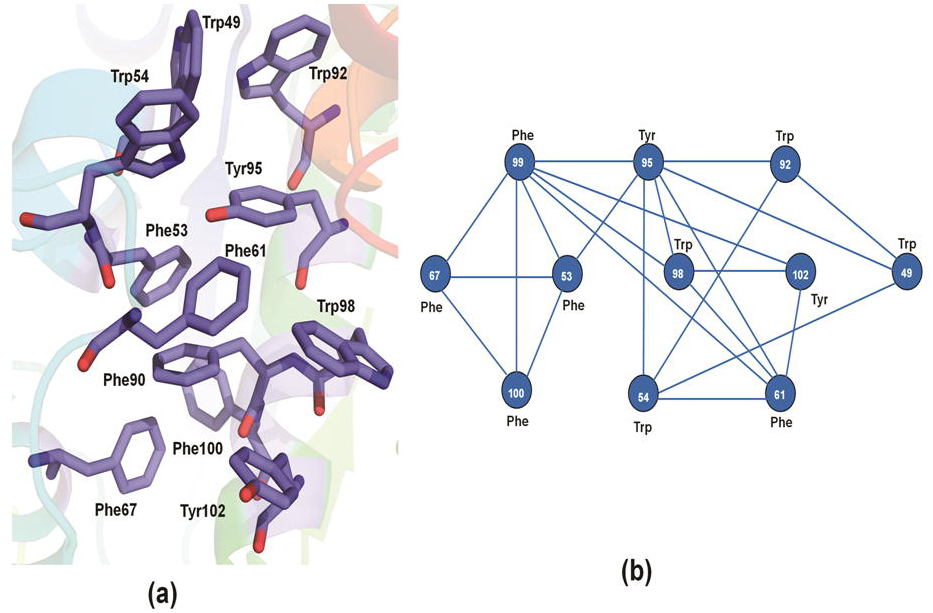
A three-residue-clique as a building block of large aromatic cluster. (a) A cluster of eleven aromatic residues of a k =3 aromatic-clique community for thermostable Rhodanese (T_m_= 70 °C) from *Thermus thermophilus HB8* (PDB ID: 1UAR) that comprises of twelve three residue cliques. (b) A schematic view of the aromatic-clique community presents in thermostable Rhodanese from *Thermus thermophilus HB8*. It may be seen that intermingling of k = 3 cliques results in higher order cliques of aromatic-aromatic interactions and emergence of k = 4 cliques is also seen (Phe53-Phe67-Phe99-Phe100; Phe53-Phe61-Tyr95-Trp98-Phe99-Tyr102).

## Discussion

In the present study, we report a systematic investigation of a set of aromatic mutant crystal structures of a thermostable xylanase from *Bacillus sp. NG-27* and provide a structural insight into the role of long-range aromatic interaction network particularly between N-terminal and C-terminal, in protein stabilization. Figure 1 provides an example of such interactions in RBSX structure in which the disruption of long-range aromatic interactions (F4A, W6A and Y343A) resulted in substantial reduction in the thermal stability of the enzyme, whereas aromatic to aromatic substitution (F4W), when the aromatic clique community is maintained, showed no alternation in protein stability (Fig. S1; Table S1). It appears that the decreased stability shown by these aromatic mutants (F4A, W6A and Y343A) resulted from a cumulative effect in the loss of aromatic interactions and reduced van der Waals interactions within N-terminal region and between N-terminus and C-terminus regions. Various studies have shown the role of interactions between termini in modulating the stability of proteins [15-18]. An *in silico* analysis of a set of two-state folding proteins showed the presence of an N-C motif (N-terminal to C-terminal contacts) and suggested its possible role in initial protein folding, native state stability and final turnover [19]. In a recent study, we have provided experimental evidence through crystal structure analysis of different extreme N-terminus aliphatic mutants of RBSX where augmenting N-terminal to C-terminal interactions is associated with enhancement of the stability of the enzyme [11]. The present results are consistent with earlier findings and exemplify the importance of interactions between N-terminus and C-terminus specifically through aromatic interactions and provide a network perspective of aromatic interactions/cluster in modulating the stability of the enzyme.

The importance of aromatic interactions has been reported previously in association with protein thermostability [6, 20-21]. A mutational study of CYP119 from *sulfolobus solfataricus*, it was shown that aromatic interaction is an important determinant of the thermostability of the protein [20]. They found that mutation of individual residues (W231A, Y250A and W281A) lowered the T_m_ of the mutants by ∼10-15 °C with respect to the wild type protein. Interestingly, in the structural analysis of the wild type protein, CYP119 (PDB ID: 1IO8), we observed that all the three mutated aromatic residues (Trp231, Tyr250 and Trp281) belong to three different long-range aromatic-clique communities Phe225-Trp231-Tyr240, Phe228-Tyr250-Phe298, and Trp4-Phe5-Trp281, revealing the importance of long-range aromatic cluster in the form of aromatic-clique communities in the structural stabilization of the protein (Fig. 4a). There have been other experimental reports of the effect of aromatic mutation/s on the structural stability of proteins. For example, the substitution of various partially exposed phenylalanines (Phe15, Phe17 and Phe27) to alanine destabilized the folded state of a cold shock protein B from *Bacillus substilis* [22]. In the analysis of the structure of the cold shock protein (PDB ID: 1CSP), we found that Phe15, Phe17 and Phe27 are involved in aromatic interaction and form an aromatic-clique community (F15-F17-F27) (Fig. 4b). Another such example comes from the protein from *Staphylococcal Nuclease* (PDB ID: 1STN) in which various systematic mutations including F34A, F34G, F76A and F76G in the core domain significantly decreased the stability of the protein [23]. We observed that both Phe34 and Phe76 belong to an aromatic-clique community composed of residues Tyr27, Phe34 and Phe76, affirming the role of long-range aromatic-clique interaction in protein stability (Fig. 4c). Thus, we can interpret our results and that of others in terms of long-range aromatic clique community and infer that a disruption of the community structure results in destabilization. In addition, the present observations reinforce a view that is emerging from a comparative analysis of crystal structures of homologous pairs of proteins from thermophilic and mesophilic organisms and show the preponderance of aromatic-clique communities in thermophilic proteins in comparison with their mesophilic counterparts (Table S3). Aromatic residues are found to participate in interaction networks and have been reported to enhance the stability of proteins, mainly via dimer or cluster formation [6-8]. However, the aromatic clusters that form a k-clique community offer an efficient clustering, and can be expected to be a more cooperative and cohesive structure for coordination and cooperation. On the other hand, a cluster formed by k-clique communities has additional advantages due to their interaction synergy and better effective interaction energy for stabilization [8]. Furthermore, the long-range aromatic-clique community can enable to bridge distant regions of the protein structure which are far apart in sequence space, providing additional stabilization to the tertiary structure of the protein [24-26]. Taken together, these results highlight an additional source of stability which could arise due to the prevalence of long-range aromatic-cliques and communities by increasing the mutual interactions within the cluster and through long-range interactions (in terms of sequence separation), connecting different parts of the protein structure.

**Fig. 4.**
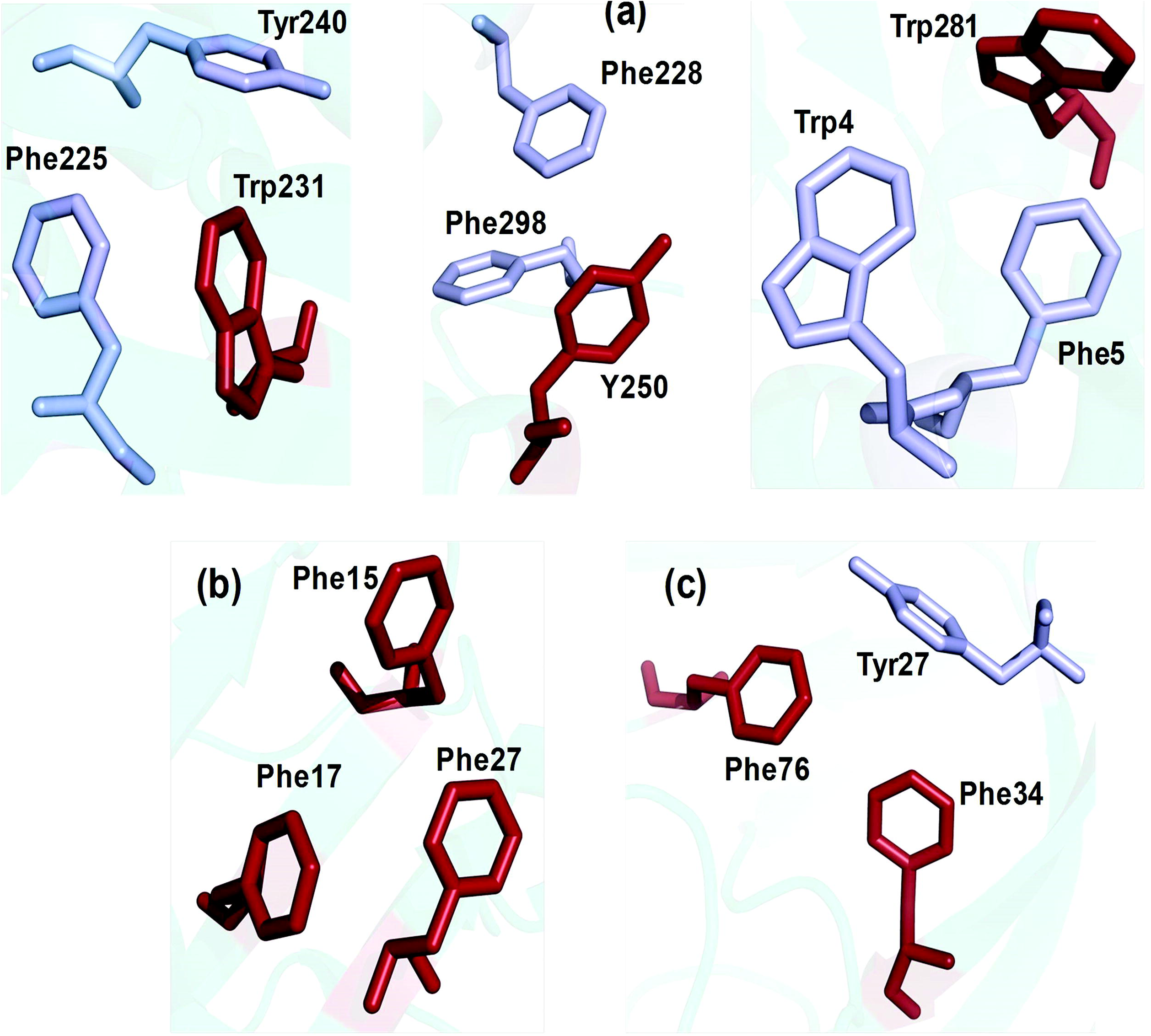
The association of long-range aromatic k-clique community with protein stability. The mutated residue/s is represented as stick in firebrick. (a) The mutations in long-range aromatic-clique communities of CYP119 from *sulfolobus solfataricus* (PDB ID: 1IO8), (b) cold shock protein from *Bacillus substilis* (PDB ID: **1CSP**) and (c) core domain of *Staphylococcal nuclease* (PDB ID: 1STN), modulate the stability of the protein.

## Conclusion

The aromatic-clique interaction (Phe4-Trp6-Tyr343) in RBSX is not directly linked to catalytic mechanisms but has a profound effect on the thermal stability of the protein. Even though individual studies of aromatic mutations affecting protein stability have been reported earlier, our works brings together such studies by a systematic analysis of crystal structures of aromatic mutants and elucidated the role of aromatic clique community, long-range interaction and N-terminal to C-terminal interaction in protein stability. The rigid and planar moieties of the aromatic residues come into close proximity in 3D space from different parts of the protein structure, providing opportunities to interact with one another, abut one another, form clique communities and bolster the 3D structure for the stabilization of protein (Fig. 3 and Fig. 4). The stabilization of 3D structure through long-range clique communities of interacting aromatic residues is present in other proteins (Fig. 4, Table S3). In the present study, based on our experimental work and that of others, we show that disruption of the aromatic community structure through mutations results in protein destabilization (Fig. 1 and Fig. 4). Community structure is one of the most relevant features of graph or network representing real systems, and has been discussed in social, economic, computer science, and biological networks [27-30]. The community analysis can provide a conceptual framework for the study of correlated motions in nucleic acids and proteins, protein-protein interactions networks, metabolic networks and propagation of allosteric signals in proteins, underlining the importance of such approach [31-35]. Furthermore, we show here that aromatic-clique community offers an interesting perspective to analyze aromatic cluster/s in protein structures and their implications for protein stability. Furthermore, we point out that a three-node aromatic clique can act as building block of large/complex network motifs in protein structure (Fig. 2, Fig. 3 and Table S3). We suggest that the strategy of mutations based on clustering of aromatic residues in the form of aromatic clique and community could be effectively applied to enhance the stability of enzymes and provide a new approach for the design of proteins for biotechnological applications.

## Materials and Methods

### Aromatic-clique definition, detection and long-range interaction community

In the present case, Phe, Tyr, and Trp are considered as aromatic residues. Histidine is not considered as an aromatic residue because of its polar and hydrophilic character and involvement in other types of interactions [8]. Two aromatic residues are defined to form an aromatic pair if the closest distance between any two atoms (except hydrogen) of the rings is < 6.0 Å and the distance between ring centroids (considered only the benzene ring for Phe and Tyr and centroid of five atoms of the pyrrole ring for Trp) is < 8.5 Å. With the aromatic pairs calculated by the above definition and criteria, communities of aromatic interactions are drawn with CFinder software [31, 36]. This software shows k-clique communities within a network according to k-values and nodes belonging to each community. A k-clique (fully connected subgraph of k-nodes) is defined as a set of k nodes (Phe, Tyr and Trp) in which each node is connected to all the other nodes. A community can be defined as a union of smaller complete subgraphs that share nodes. In general, a k-clique community is defined as a union of all k-cliques that can be reached from each other through a series of adjacent k-cliques. The higher order cliques are rare in protein interaction network because the edges between nodes are determined by distances and there is a minimal possibility to form a more complex network. Hence, we have focused on only k=3 k-cliques communities of aromatic interaction for the study to find an aromatic cluster. Furthermore, we have filtered out all the k=3 k-cliques communities based on sequence distance by the clique forming residues. We considered only those k=3 k-clique communities for which the relative sequence distance between any two nodes in k-clique community is greater than 10, thereby focusing on long-range interactions [25].

### Data set of Thermophilic GH10 xylanases

The crystal structures of thermophilic GH10 xylanases were obtained from the CAZY database (http://www.cazy.org) and downloaded from the Protein Data Bank (PDB) database (http://www.rcsb.org/pdb/home/home.do). In total, sixteen GH10 xylanases from different thermophilic organisms were selected and analyzed for the study. First, aromatic interactions were identified and subsequently aromatic-clique was detected based on the definition of k-clique communities as described above.

### Aromatic-clique and community: Beyond GH10 xylanases

We have constructed a data set of 57 homologous pairs of proteins from thermophilic and mesophilic organisms from different protein families for the analysis of aromatic-clique interactions. Only 3D structures obtained from X-ray crystallographic method are used. Structures determined using nuclear magnetic resonance (NMR) methods and modeled structures have been excluded from the dataset. Further, the dataset was filtered such that upon superposition of corresponding C_α_-atoms, the root mean square deviation (RMSD) between the thermophilic and their mesophilic homologous structure was not more than 5Å and chain lengths differ by not more than 30 amino acid residues. This filtering of the dataset ensures that the homologous proteins that were paired are structurally similar and the structural difference between the thermophilic proteins and their corresponding mesophilic homologs was due to the difference in stability but not an evolutionary artifact [37].

### Structural alignment

The molecular structures of thermophilic and their mesophilic homologs were superimposed using the ‘Superpose’ program from the CCP4 package [38]. Superpose provides a structure-based sequence alignment and was used to compare the topologically equivalent residues in the pair of homologous thermophilic and mesophilic proteins.

## Acknowledgements

P. Mahanta thanks UGC-CSIR (India) and Indian Institute of Science, Bangalore for financial support. X-ray data for F4A, F4W and Y343A crystals were collected at the X-ray Facility for Structural Biology at the Molecular Biophysics Unit, Indian Institute of Science, Bangalore, India. We thank beamline staff at BM14 of the European Synchrotron Radiation Facility (ESRF), Grenoble for providing access to collect the diffraction data of W6A mutant. The authors thank Krishan Kumar who purified the protein samples of F4A. S. Ramakumar thanks UGC (India) for an emeritus fellowship.

## Author contributions

P. Mahanta, V.S. Reddy and S. Ramakumar designed the experiments. P. Mahanta, A. Bhardwaj performed the experiment. P. Mahanta crystallized, data collected, and solved the structures. P. Mahanta carried out the computational analysis. All authors wrote the manuscript and approved the manuscript.

